# Operative and technical modifications to the Coriolis^®^ µ air sampler that improve sample recovery and biosafety during microbiological air sampling

**DOI:** 10.1101/2020.02.11.943662

**Authors:** Nuno Rufino de Sousa, Lei Shen, David Silcott, Charles J. Call, Antonio Gigliotti Rothfuchs

**Author notes:** Address correspondence to Antonio Gigliotti Rothfuchs, MTC, Karolinska Institutet, Biomedicum, Solnavägen 9, SE-171 77 Stockholm, Sweden. These authors contributed equally to this work (Nuno Rufino de Sousa, Lei Shen). Shanghai Pulmonary Hospital, Tongji University School of Medicine, Shanghai, China (Lei Shen).

## Abstract

Detecting infectious aerosols is central for gauging and countering airborne threats. In this regard the Coriolis^**®**^ µ cyclonic air sampler is a practical, commercial collector that can be used with various analysis methods to monitor pathogens in air. However, information on how to operate this unit under optimal sampling and biosafety conditions is limited. We investigated Coriolis performance in aerosol dispersal experiments with polystyrene microspheres and *Bacillus globigii* spores. We report inconsistent sample recovery from the collector cone due to loss of material when sampling continuously for more than 30 min. Introducing a new collector cone every 10 min improved this shortcoming. Moreover, we found that several surfaces on the device become contaminated during sampling. Adapting a HEPA-filter system to the Coriolis prevented contamination without altering collection efficiency or tactical deployment. A Coriolis modified with these operative and technical improvements was used to collect aerosols carrying microspheres released inside a Biosafety Level-3 laboratory during simulations of microbiological spills and aerosol dispersals. In summary, we provide operative and technical solutions to the Coriolis that optimize microbiological air sampling and improve biosafety.

## INTRODUCTION

*Mycobacterium tuberculosis*, measles virus, influenza virus and other highly contagious human pathogens transmit through air, either by aerosol or droplet transmission (Riley et al., 1978; Bloch et al., 1985; Remington et al., 1985; Fennelly et al., 2004; Yang et al., 2011; Cowling et al., 2013; Patterson et al., 2017). These airborne pathogens pose a heavy burden on society by incurring a spectrum of outcomes ranging from death to morbidity to absence from work due to sickness. Airborne microbes are of particular concern in enclosed, crowded environments, where occupants are readily exposed to respired air and thus at risk of inhaling infectious bioaerosols carrying viruses, bacteria or fungi. This is well-recognized during infection with *M. tuberculosis* where congregate settings such as prisons, homeless shelters, slums and refugee camps are recognized hotspots of transmission (WHO, 2009).

Bioaerosols are produced during coughing or sneezing (Nicas et al., 2005; Yang et al., 2007; Fernstrom and Goldblatt, 2013) and at lower concentrations during talking or breathing (Diffey, 2011; Fernstrom and Goldblatt, 2013). The concentration of bioaerosols in a given indoor setting depends on occupants’ consumption of oxygen, respiratory quotient and physical activities (Persily, 1997; Emmerich and Persily, 2001), as well as physical factors of the indoor environment, such as ventilation rate, number of occupants and room volume (Persily, 1997; Emmerich and Persily, 2001; Lygizos et al., 2013). Since individuals spend the majority of the working hours of the day indoors (Diffey, 2011), enclosed environments pose a general risk for acquiring airborne infections.

Many advancements have been made in our understanding of aerosol formation and dispersion, on the risks of exposure and on ways to interfere with transmission of airborne pathogens. In this context, the ability to monitor pathogens in air is an important investment for gauging and controlling infectious disease in society. Microbiological air-sampling tools enable detection of pathogens in air and as such improve our position to counter airborne threats through capacity-building, infection control measures. In particular, there is an outstanding need to monitor pathogens in air in critical infrastructure such as government buildings, hospitals, mass transit and airports, during manufacturing in clean-rooms and “ready-to-eat” food preparation, to name but a few.

The Coriolis^®^ µ (Bertin Instruments, Montigny-le-Bretonneux, France) is a state-of-the-art, high-volume air sampler that collects airborne particles into liquid through cyclonic-air sampling. The unit is high-cost and energy-hungry but has tactical capacity and produces a sample that is compatible with many different analytical methods. The Coriolis has been used in a variety of air-sampling applications, including collection of chemical compounds (Caygill et al., 2013), toxins (Viegas et al., 2012) and microbial contaminants in the food industry (Verreault et al., 2011; Viegas et al., 2014). It has also been used for surveillance of airborne pathogens in healthcare facilities (Le Gal et al., 2015; Montagna et al., 2017; Alsved et al., 2019; Montagna et al., 2019). It is known that during sampling the collecting cone loses a considerable amount of liquid, raising the possibility of inner contamination of the device and that longer sampling intervals may incur unintentional re-aerosolization and exposure of the microorganisms sampled for analysis, a situation of especial concern in the latter cases.

Despite its use in different microbiological air-sampling applications, investigation on actual Coriolis performance and biosafety concerns during operation have not been thoroughly addressed. Herein we have evaluated the Coriolis in a series of aerosol collection experiments with microspheres and bacterial spores under controlled laboratory conditions. We report on a sampling protocol to maximize sample recovery from the unit and a HEPA-filter adaptation to reduce unintentional contamination of device parts which occurs as a consequence of re-aerosolization of collected material during sampling. We demonstrate the use of the modified Coriolis in the detection of aerosols generated during a simulated laboratory spill and aerosol dispersal.

## METHODS

### Bacteria and fluorescent beads

Lyophilized endospores of *Bacillus atrophaeus* var. *globigii* (*Bg*)(from ECBC Pine Bluff Arsenal Laboratories, US Army, originally given to D. Silcott) were resuspended in sterile deionized (DI) water. Quantification of *Bg* Colony-forming units (CFUs) in stocks and aerosol samples was determined by culture on LB Miller agar (Sigma-Aldrich) for one day at 37 °C. Stock solutions of *Bg* were diluted to 1×10^9^ CFUs/mL and stored at 4 °C until further use. Stock solutions of 1 µm, yellow-green (505/515) fluorescent, polystyrene FluoSpheres™ (ThermoFisher) were also prepared in DI water at 1×10^9^ beads/mL, stored at 4 °C and protected from light until further use.

### Aerosol dispersal experiments in a containment chamber

Contained aerosol dispersal experiments were performed inside a large, airtight, flexible PVC enclosure mounted on a metal-support frame (Solo Containment, UK). The enclosure measures 270 cm (L) x165 cm (W) × 255 cm (H) with an inner volume of 9.3 m^3^. It was assembled and kept inside a Biosafety Level (BSL)-2 laboratory. Particles were purged from the chamber before the start of experiments by drawing air into the enclosure through a HEPA filter using an Attix 30 industrial-grade vaccuum cleaner (Nilfisk, Sweden). FluoSpheres or *Bg* were aerosolized into the chamber using a 4-jet Baustein atomizing module (BLAM) nebulizer (CH Technologies. USA) operated in multi-pass mode and producing quasi-monodisperse aerosols with a particle-size diameter range of 0.7-2.5 μm. Aerosolization was performed for 1 min. Previous tests established a steady concentration of particles in the enclosure 8 min after completing the aerosolization cycle on the BLAM (**Fig. 2C**) and (data not shown); air sampling was therefore routinely initiated 8 min after finishing the aerosolization cycle. In certain experiments, a Lighthouse Handheld 3016 particle counter (Lighthouse Worldwide Solutions, USA) or an IBAC fluorescent particle counter (FLIR Systems, USA) were used to measure the decay of aerosolized FluoSpheres inside the enclosure.

**Figure 1.**
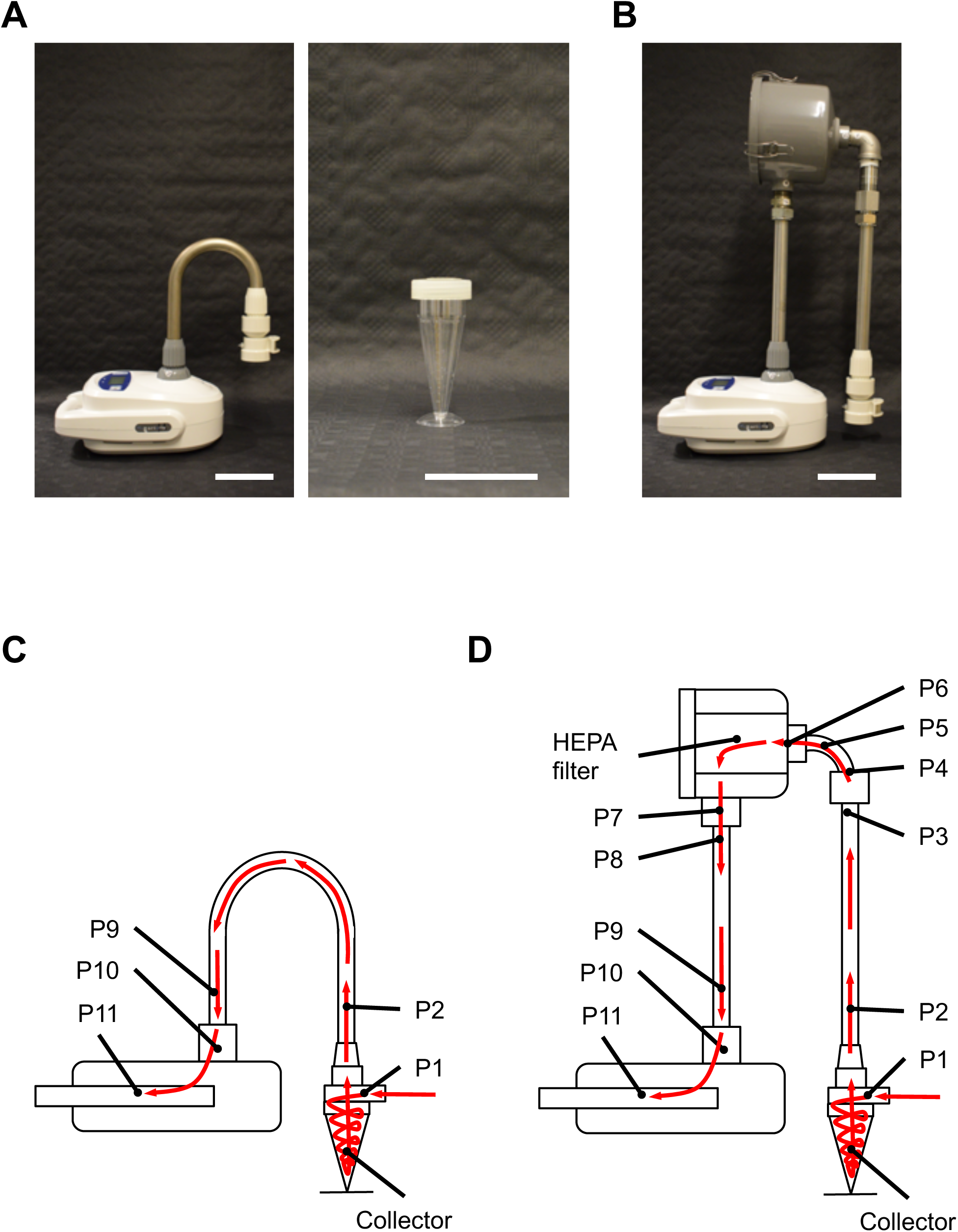
Coriolis® µ and HEPA-filter adaptation. **A**, Coriolis as supplied by vendor with accompanying parts (left panel) including collector cone (right-center panel). **B**, Coriolis after incorporation of customized HEPA-filter adaptation with parts presented in **Supplementary Fig. 1**. Scale bars depicting 10 cm. **C-D**, Cartoon of Coriolis internal components without (**C)** and with HEPA filter (**D**). Arrows in red showing direction of sampled air through the device. P1-P11 denotes locations from which swabs were obtained for regrowth of *Bg* in experiments presented in **Fig. 4A-B**.

**Figure 2.**
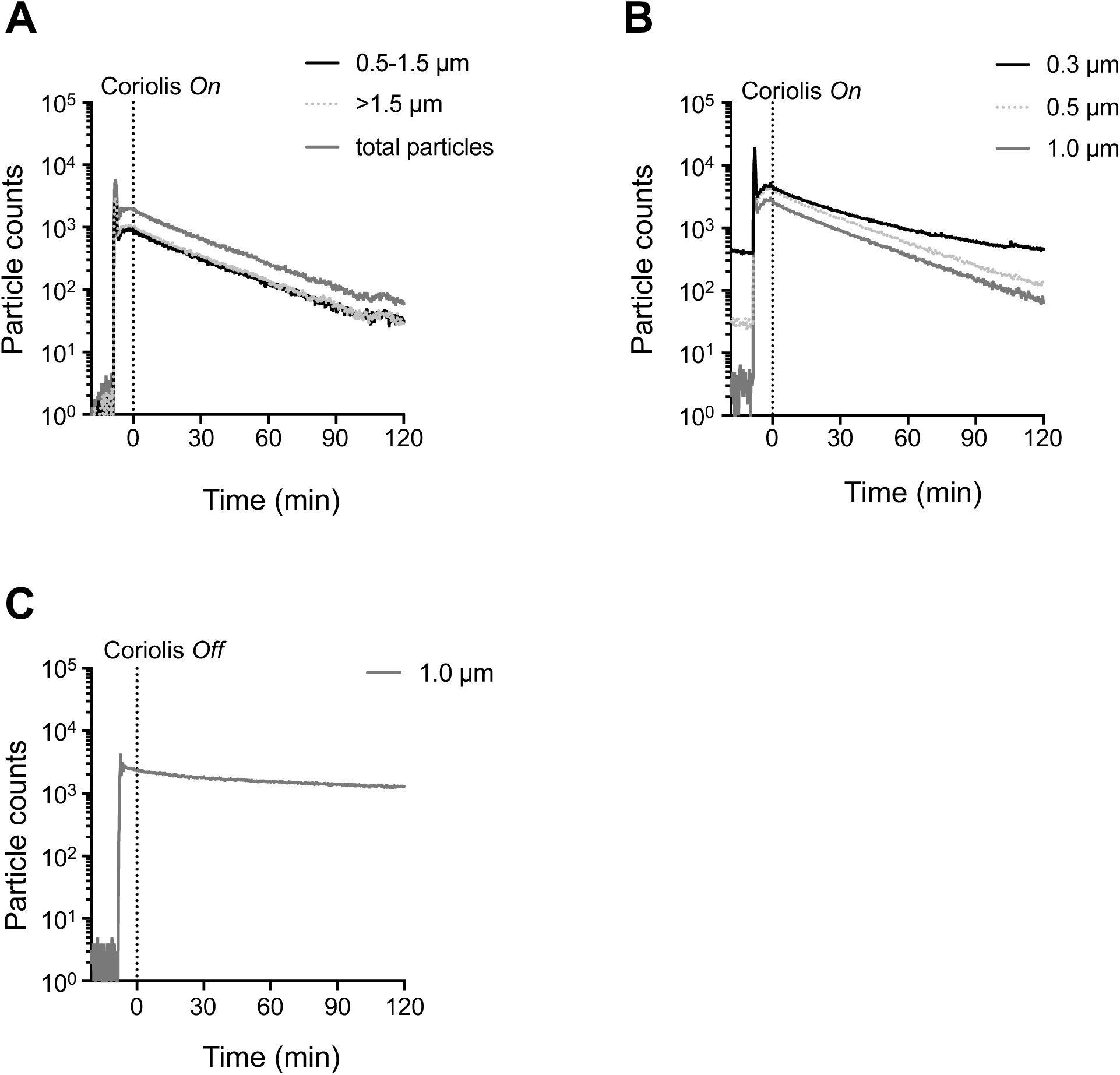
Clearance of aerosolized microspheres from air during Coriolis sampling. **A-C**, FluoSpheres (1 µm, 1×10^9^) were aerosolized inside the aerosol chamber. Coriolis was turned *on* or left *off* (dashed line) and particle counters used to record microspheres in the air. **A**, Fluorescent particle counts recorded on an IBAC sensor with Coriolis *on*. **B-C**, Particle counts recorded on a Lighthouse particle counter with Coriolis *on* (**B**) and at steady state with Coriolis *off* (**C**).

### Coriolis® µ function and HEPA-filter adaption

A Coriolis^®^ µ cyclonic air sampler (Bertin Instruments, Montigny-le-Bretonneux, France) (**Fig. 1A**) was used to collect aerosolized microparticles in controlled, aerosol dispersal experiments. A bespoke solution for integrating a HEPA-filter system to the Coriolis was conceived and assembled as illustrated (**Fig. 1B**) with necessary filter component, stainless-steel metal tubing and other fittings (**Supplementary Fig. 1**) (all MacMaster-Carr, USA). Air flow measurements were made on the Coriolis with a TSI 4040 mass flow meter (TSI, USA) connected to the blower inlet of the Coriolis using thin-wall latex tubing, stretched such that a tight seal was obtained on both the air-flow meter and the Coriolis inlet. Measurements were made at the manufacturer-specified flow rate setting of 300 L_air_/min. The flow rate measured was 298 L_air_/min without a collector cone (**Fig. 1A**) connected to the device. With a collector cone loaded with 15 mL connected to the Coriolis the flow rate measured was 270 L_air_/min. With the HEPA filter attached to the unit and loaded with a collector cone containing 15 mL, the flow rate at the inlet was 250 L_air_/min.

For experiments performed in the containment chamber, the Coriolis was placed on a table 30 cm away from the BLAM aerosol port with the collector part of the unit approximately 100 cm from the ground. As per manufacturer’s recommendation, the collector cones were filled with 15 mL DI water and kept sealed. After completion of the aerosolization cycle, a collector cone was unsealed, loaded onto the Coriolis and the sampler operated at the manufacturer-designated flow rate of 300 L_air_/min. Collection on the Coriolis was performed using the same cone for the entire sampling duration and referred to as *standard sampling*. In this setting an injection port was used to manually replenish the collector cone back to 15 mL of DI water after every 10 min of sampling. Alternatively, a new collector cone containing 15 mL of DI water was replaced after every 10 min of sampling, referred to as *cumulative sampling*. At the end of experiments, the enclosure was decontaminated with 35% hydrogen peroxide vapor using a BQ-50 unit (Bioquell, UK). Before starting this procedure, the removable metal piping was disassembled from the body of the Coriolis and the Coriolis operated in decontamination mode. Metal piping was cleaned in mild detergent and autoclaved.

### Spill and aerosol dispersal experiments

Laboratory spill and aerosol dispersal simulations were performed inside a suite of the BSL-3 facility at Biomedicum, Karolinska Institutet, Solna, Sweden. The suite dimensions measure 1000 cm (L) × 300 cm (W) × 270 cm (H). The facility operates at approximately 20-27 air-changes-per-hour (ACH) depending on the number of microbiological safety cabinets in simultaneous use. Safety cabinets in the suite were inactive during our experiments, lowering forced ventilation parameters to about 18 ACH. Our experiments were done before the BSL-3 was opened to users and pathogens introduced to the facility. For spill simulations, the Coriolis was placed approximately 10 cm from the planned spill site and rested either on top of a working bench (90 cm from the ground) or just above the ground (30 cm). A large microbiological spill was simulated by tumbling a container carrying 0.5 L of FluoSpheres (2×10^6^ beads/mL in DI water, 1×10^9^ beads in total) from the same working bench resting the Coriolis. For aerosol dispersal experiments, the Coriolis was placed on the working bench 90 cm from the ground. The Coriolis was located 900 cm in front of the BLAM, which was supported on a tripod 100 cm from the ground. The BLAM was loaded with FluoSpheres (1×10^9^ beads/mL). Aerosolization was performed for 1 min, releasing a maximum of 1×10^9^ microspheres into the room. In both simulations, air sampling was performed on the Coriolis with accompanying HEPA-filter modification (**Fig. 1B**) for 1 hr using the cumulative sampling method described above. Researchers donned full-body Tyvek® 500 Labo overalls, 3M® FFP3D respirators, safety goggles and nitrile gloves (all VWR), providing all-around contact protection from aerosol exposure. At the end of experiments, working surfaces were cleaned with mild detergent and the entire BSL-3 suite was decontaminated with 35 % hydrogen peroxide vapor using a BQ-50 unit.

### Swabbing and extraction of swab samples from Coriolis parts

At the end of a collection cycle on the Coriolis, the aerosol chamber was purged from airborne particles as described above. The Coriolis was then swabbed using sterile cotton swabs (VWR) at the designated points P1-P11 in **Fig. 1C-D**. Sample was extracted from swabs by breaking the cotton-end of the swab into a sterile microcentrifuge tube. 0.5 mL PBS-0.05% Tween-80 was added to the tube, the tube was sealed, incubated at room temperature for 2 min and vortexed for 1 min. The cotton-end of the swab was then aseptically removed from the tube using a pair of sterile tweezers.

### Calculation of particle clearance rate

The particle clearance rate for aerosolized FluoSpheres in the aerosol chamber was obtained by dividing the inner volume of the chamber (9300 L) by the amount of time needed to reach a 4-Log reduction of the starting material of FluoSpheres aerosolized in the chamber (*i.e.* 99.99% clearance). The amount of time needed to reach a 4-Log reduction was calculated by fitting a one-phase decay curve to the particle counts measured in the chamber over time using designated particle counters (IBAC sensor and Lighthouse particle counter). The particle clearance rate for the Coriolis was expressed in L_air_/min.

### Flow cytometric quantification of FluoSpheres

FluoSpheres collected on the Coriolis were quantified on a flow cytometer using CountBright™ Absolute Counting Beads (ThermoFisher). Briefly, a defined number of CountBright beads were added to 1 mL of sample, samples were acquired on a FACS Calibur (BD Biosciences) and analyzed on FlowJo (BD Biosciences). The number of FluoSpheres in the sample was calculated with CountBright™ beads according to the instructions of the manufacturer (ThermoFisher).

### Statistical analyses

The significance of differences in data group means was analyzed using Student’s *t* test or Anova where appropriate, with a cut-off of p< 0.05.

## RESULTS

### Rapid decay of aerosolized particles from air during Coriolis sampling

A robust air sampler must be able to effectively collect particles from air while also generating a sample that is amenable to downstream analysis. For the Coriolis (**Fig. 1A**), we investigated particle collection indirectly by measuring the unit’s particle clearance rate, *i.e.* the rate at which the Coriolis removes airborne particles from air. Fluorescent polystyrene (1 µm) FluoSpheres were aerosolized in the aerosol chamber in the presence of the Coriolis and airborne particles measured in real-time with an IBAC sensor and a Lighthouse particle counter, respectively. When the Coriolis was turned *on* the number of airborne particles recorded in the chamber began to steadily decay and decreased about 1.5 orders of magnitude in 2 hrs (**Fig. 2A-B**). On the contrary, particle decay was not observed when the Coriolis was left *off* (**Fig. 2C**). With this information the effective clearance rate for the aerosolized microspheres was calculated to approximately 35 L_air_/min. This number was reached with the help of either particle counter. Additional size-discriminating particle analysis with the Lighthouse particle counter showed that the Coriolis struggled with particles smaller than 0.3 µm (**Fig. 2B**), in line with technical specifications reported by the manufacturer (Bertin Instruments).

### A cumulative sampling method that improves sample recovery during Coriolis air sampling

Pathogen numbers in aerosols are limited. Thus, any protocol that improves collection or minimizes material loss adds value to the use of that collector during microbiological air sampling. We aerosolized FluoSpheres in the aerosol chamber and investigated their collection on the Coriolis over time to see if this process could be improved. Collection was performed every 10 min for a total of 2 hrs. The sample volume on the collector cone was manually replenished to 15 mL every 10 min as recommended by the manufacturer. Indeed, although the Coriolis is reported by the manufacturer to be able to collect material for up to 6 hrs, we found that sample recovery was inconsistent after 30 min of sampling (**Fig. 3A**). We hypothesized that the collected material might be escaping the collector cone over time as a consequence of cyclonic sampling. Hence, we modified the sampling procedure by replacing the collector cone after every 10-min cycle with a new one; analyzing each cone individually and cumulatively adding the quantification obtained from each time-point to the detection curve. At the end of the 2 hr-sampling interval we found that this *cumulative sampling* protocol lead to a 50% improvement of collection compared to the standard, manufacturer-recommended sampling with manual liquid replenishment (**Fig. 3A**). Analysis of material recovered from individual time-points during cumulative sampling showed that approximately 95% of the FluoSpheres recovered during the 2hr-sampling interval were collected within the first 60 min (**Fig. 3B**). Still, an appreciable amount of microspheres were collected by the Coriolis during the remaining 60 min of sampling. Overall, our observations suggest that the cumulative sampling protocol may be especially useful for long-term sampling applications.

**Figure 3.**
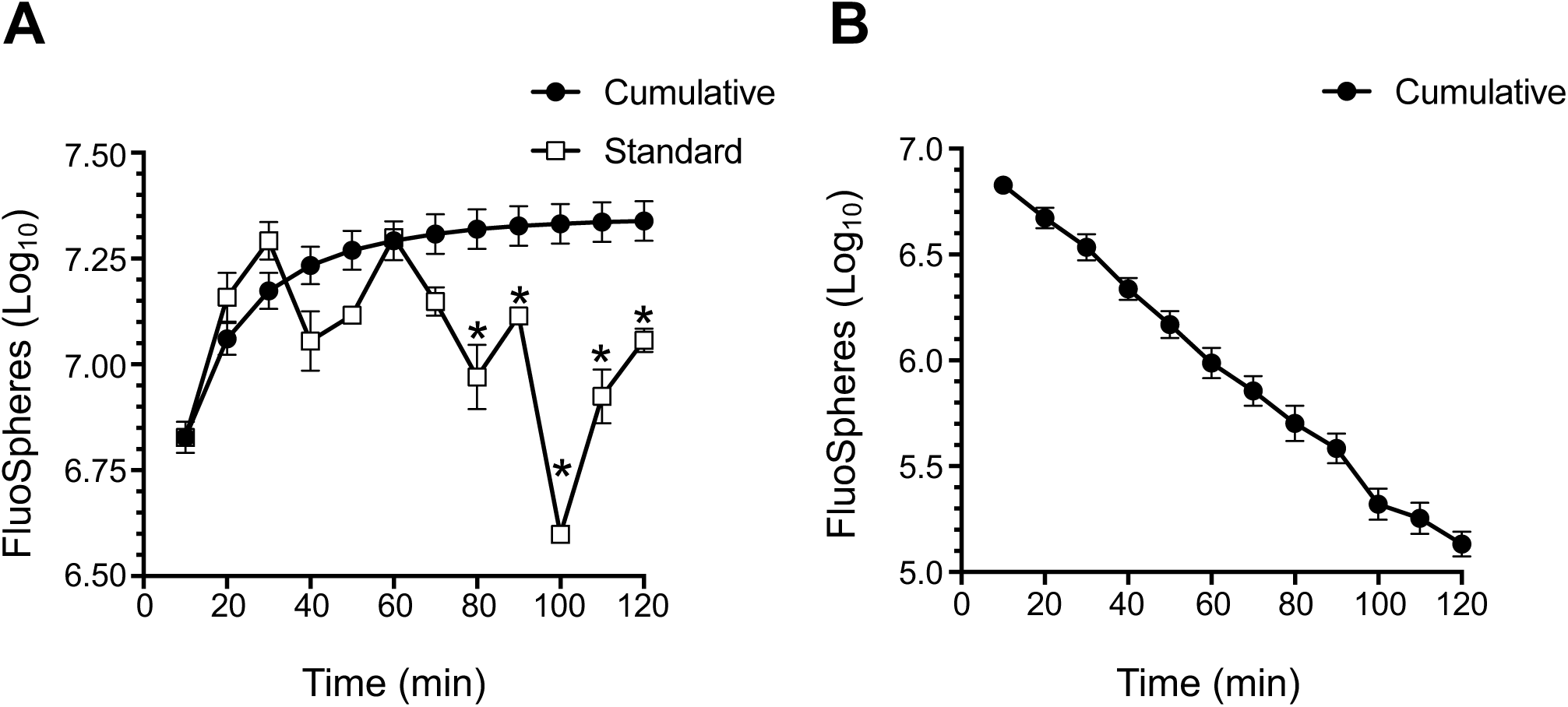
Cumulative sampling improves sample recovery from Coriolis during prolonged air sampling. FluoSpheres (1 µm, 1×10^9^) were aerosolized in the aerosol chamber as in **Fig. 2** and collected on the Coriolis (**Fig. 1A). A**, FluoSpheres were aerosolized in the chamber and Coriolis sampling performed continuously (Standard) or according to the *cumulative sampling* method described in the text (Cumulative). FluoSpheres collected on the Coriolis were quantified by flow cytometry. Each time point represents a separate aerosol release. For *cumulative sampling*, the number of recovered FluoSpheres from a time point is summed to the counts obtained from previous time points and graphed, meaning that each data point is showing the cumulative value of the collection up to that time point. **B**, alternative representation of data following *cumulative sampling* in (**A**) showing recovered material from each time point, *i.e.*, individual collector cones. Data from 5 experimental repeats shown. Error bars show standard error of the mean. * denotes statistically significant differences between Standard and Cumulative sampling methods.

### Bioaerosol contamination of device parts during sampling

Given that sample recovery decreased with increasing sampling time, we asked whether sampled material redistributed to other parts of the Coriolis during operation. To investigate this, we decided to swab various surfaces of the Coriolis after air sampling to see if any parts other than the collector cone became positive after collection. We chose to aerosolize *Bacillus globigii* (*Bg*) spores, the Anthrax simulant, since we have successfully extracted *Bg* by surface swabs in the past (Rufino de Sousa et al., 2020). Thus, we aerosolized *Bg* spores in the chamber and used the Coriolis to collect *Bg* bioaerosols. We then investigated regrowth of *Bg* from different parts of the Coriolis, more specifically, the collector cone and parts P1-P11 according to the schematics in (**Fig. 1C**). We found that many surfaces on the device exposed to air flow were contaminated with *Bg*. Substantial regrowth of *Bg* was obtained from the headpiece inlet (P1) where air enters the device (**Fig. 4A**). Bacilli were also readily detected at the initial tubing after the collector cone (P2) and importantly, at the air outlet (P11) (**Fig. 4A**), suggesting that bacteria may become deposited on the unit’s fan as well. This raises overall concerns regarding user safety, deposition of contaminants on the fan over time and the validity of samples analyzed on the unit.

**Figure 4.**
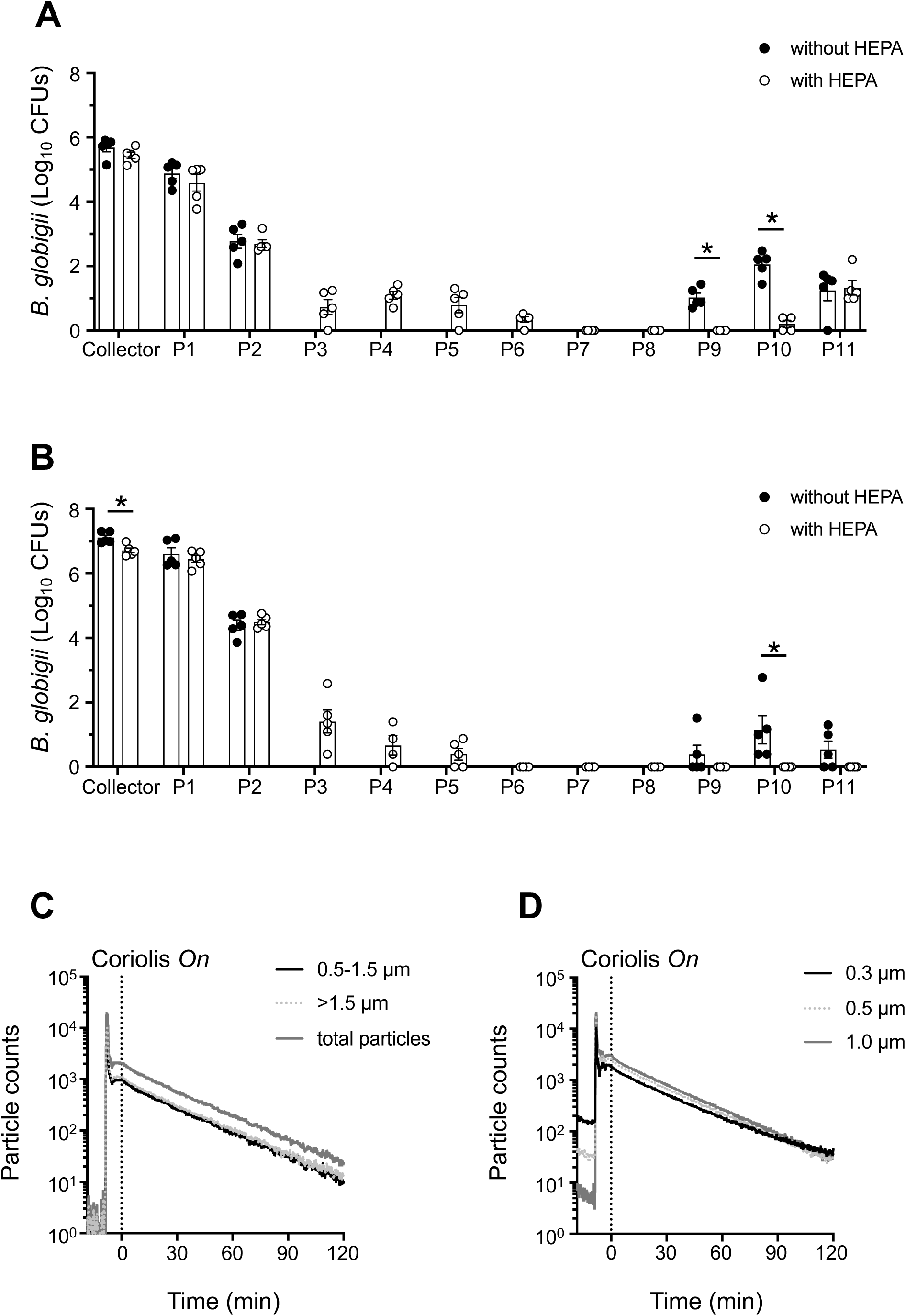
Regrowth of *Bg* from different parts of the Coriolis before and after introduction of the HEPA-filter system. **A**, *Bg* spores (1×10^8^ CFUs) were aerosolized inside the aerosol chamber and sampled for 1 hr on the Coriolis without (**Fig. 1A**) or with the HEPA-filter adaptation (**Fig. 1B**). Surface swabs were obtained from positions P1-P11 on the Coriolis (**Fig. 1C-D)**. Regrowth of *Bg* from surface swabs and the collector cone was quantified on LB agar and graphed as total number of CFUs. **B**, *Bg* spores (1×10^8^ CFUs) were loaded directly into the collector cone, the Coriolis was turned *on* for 1 hr and *Bg* regrowth investigated as in (**A**). **C-D**, FluoSpheres (1 µm, 1×10^9^) were aerosolized inside the aerosol chamber. Coriolis with the HEPA modification was turned *on* (dashed line) while particle counters were used to record airborne microspheres in the air. **C**, Fluorescent particle counts recorded on an IBAC sensor. **D**, Particle counts recorded on a Lighthouse particle counter. Data from 5 experimental repeats shown. Dots represent individual measurements. Bar graphs depict average of CFUs obtained. Error bars show standard error of the mean. * denotes statistically significant differences between Coriolis with and without the HEPA-filter modification.

In an attempt to improve device function and tactical deployment, we tailored and adapted a HEPA-filter system to the Coriolis (**Fig. 1B and D**)(**Supplementary Fig. 1**) and repeated *Bg* aerosol sampling. The HEPA filter significantly reduced *Bg* contamination from the piping (P7-P10) going into the body of the Coriolis (**Fig. 4A**). Spiking the collector cone with *Bg* confirmed that at least part of this contamination originated from re-aerosolization of bacilli in the collector cone, as the pattern of *Bg* deposition on the Coriolis was similar after aerosol dispersal (**Fig. 4A**) and spiking (**Fig. 4B**). Reducing buffer volume on the collector cone from 15 mL to 5 mL produced the same result (data not shown), suggesting that re-aerosolization was independent of the buffer volume in the cone. Importantly, there was no bacillary regrowth from the air outlet (P11) of the HEPA-modified Coriolis when the collector cone was spiked with *Bg* (**Fig. 4B**), indicating that the fan was protected from contamination in the presence of the HEPA filter. *Bg* could be detected on the outlet (P11) of the HEPA-modified unit after aerosol sampling (**Fig. 4A**). Since the HEPA filter prevents access to the outlet (P11) during spiking, detection here must be due to the high concentration of *Bg* aerosols inside the chamber, promoting deposition of bacilli onto the outside surface of the outlet, rather than a contamination coming from the inside the unit. Lastly, introducing the HEPA filter did not negatively impact on the effective clearance rate of the Coriolis (**Fig. 4C-D**).

### Use of the modified Coriolis to collect aerosols generated during a spill and aerosol dispersal

Despite biosafety and other regulatory precautions in place, the research laboratory remains an indoor environment where infections are acquired, albeit unintentionally, due to accidental exposure (Sulkin and Pike, 1951; Pike et al., 1965; Pike, 1976). With this in mind we sought to use the Coriolis with the above operative and technical modifications to investigate aerosols generated during simulated microbiological accidents in a laboratory work place. We simulated spills and aerosol dispersals in a functional BSL-3 infrastructure before it was opened to users.

To simulate a large microbiological spill, we dropped a container with 0.5 L of DI water carrying a total of 1×10^9^ (1 µm) FluoSpheres over the edge of a designated working surface in the BSL-3. A particle counter was used to record particle dispersal from the spill. Airborne FluoSpheres where collected on the Coriolis and quantified by flow cytometry. The impaction of liquid on the ground was accompanied by a detectable peak of 0.3, 0.5 and 1 µm particles close to the ground (**Fig. 5A**). To our surprise, the number of particles generated by this large liquid impaction were only marginally above baseline-particle counts in the room. Levels returned to steady-state about 10 min after the spill and were altogether undetectable when measured at the height of the working bench from which the spill was generated (**Fig. 5A**). Despite the generation of few airborne particles from this simulation, it was nevertheless possible to use the Coriolis in conjunction with flow cytometry to detect FluoSpheres aerosolized from the spill (**Fig. 5B**).

**Figure 5.**
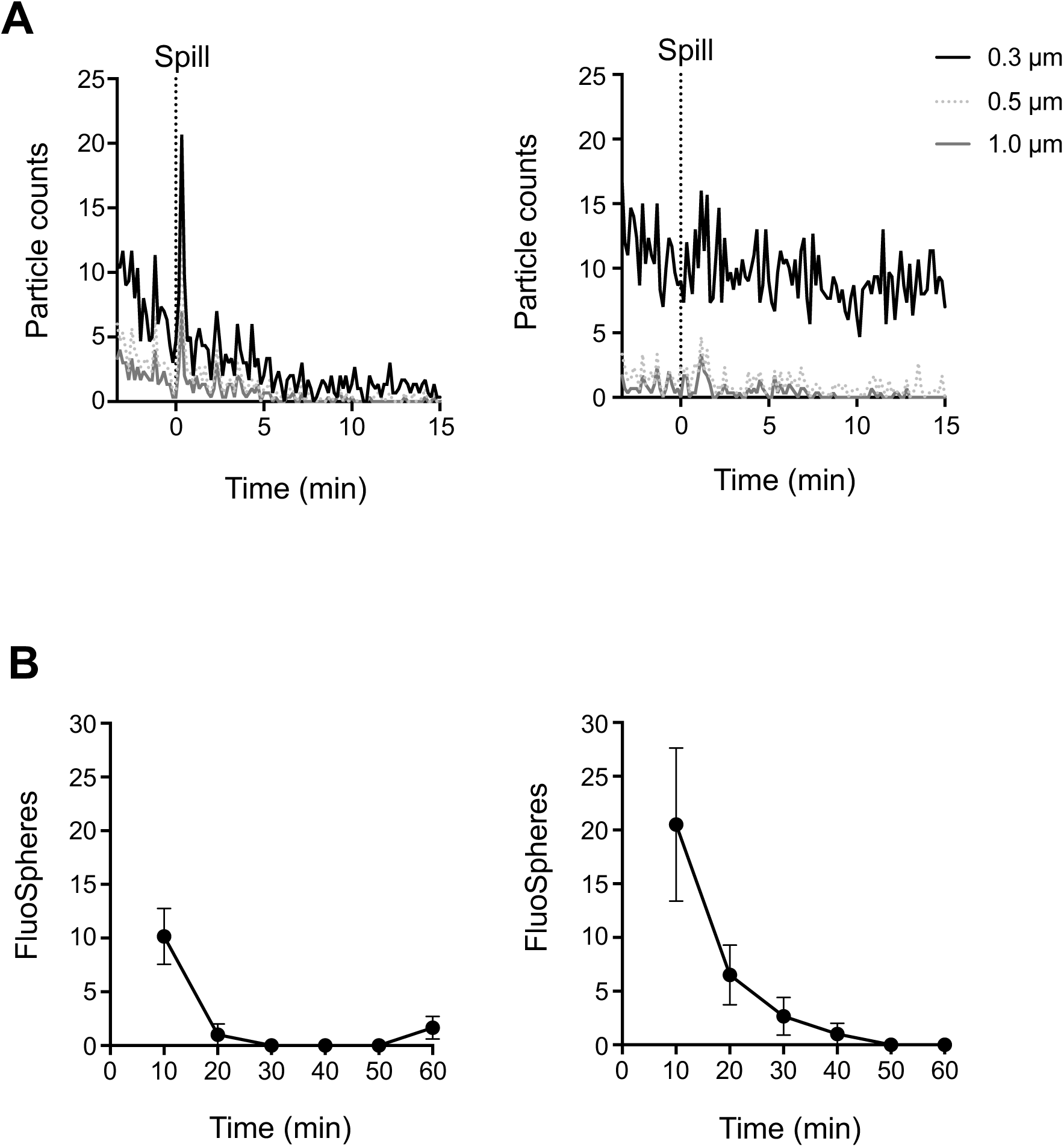
Simulation of a microbiological spill inside a BSL-3 laboratory. **A**, 0.5 L of DI water containing a total of 1×10^9^ FluoSpheres was decanted from a working surface 90 cm from the ground. A Lighthouse particle counter was used to record airborne particles at 30 cm from the ground, *i.e.* as close to the ground as possible (left panel), or 90 cm from the ground, on the working surface (right panel). **B**, The same spill was repeated and Coriolis air sampling performed with a HEPA-modified unit using the cumulative sampling method. FluoSpheres collected on the Coriolis were quantified by flow cytometry. Detection of FluoSpheres at 30 cm (left panel) or 90 cm (right panel) from the ground. Data from 5 experimental repeats shown. Error bars depict standard error of the mean. Dashed line indicates time of spill.

Next, we tested Coriolis sampling during aerosol dispersal of the same amount of FluoSpheres but on a BLAM aerosol generator. A similar peak with albeit much higher particle counts was observed concomitant with the aerosolization of these microspheres on the BLAM (**Fig. 6A**). In line, significant numbers of FluoSpheres were detected by Coriolis air sampling (**Fig. 6B**) and followed the general decay of particles in the BSL-3 suite due to forced ventilation. When central ventilation in the suite was intentionally turned *off* and the experiment repeated, airborne particle counts were increased further and remained elevated for the duration of the experiment without returning to baseline (**Fig. 6C**). Consistent with elevated and steady detection of particles in the room when central ventilation was inactivated, the Coriolis collected an elevated, steady number of FluoSpheres in the air (**Fig. 6D**). Overall, these spill and aerosol dispersal experiments reinforce the capacity of the Coriolis to enable detection of aerosolized microparticles from settings where these particles are present in not only high but also low amounts.

**Figure 6.**
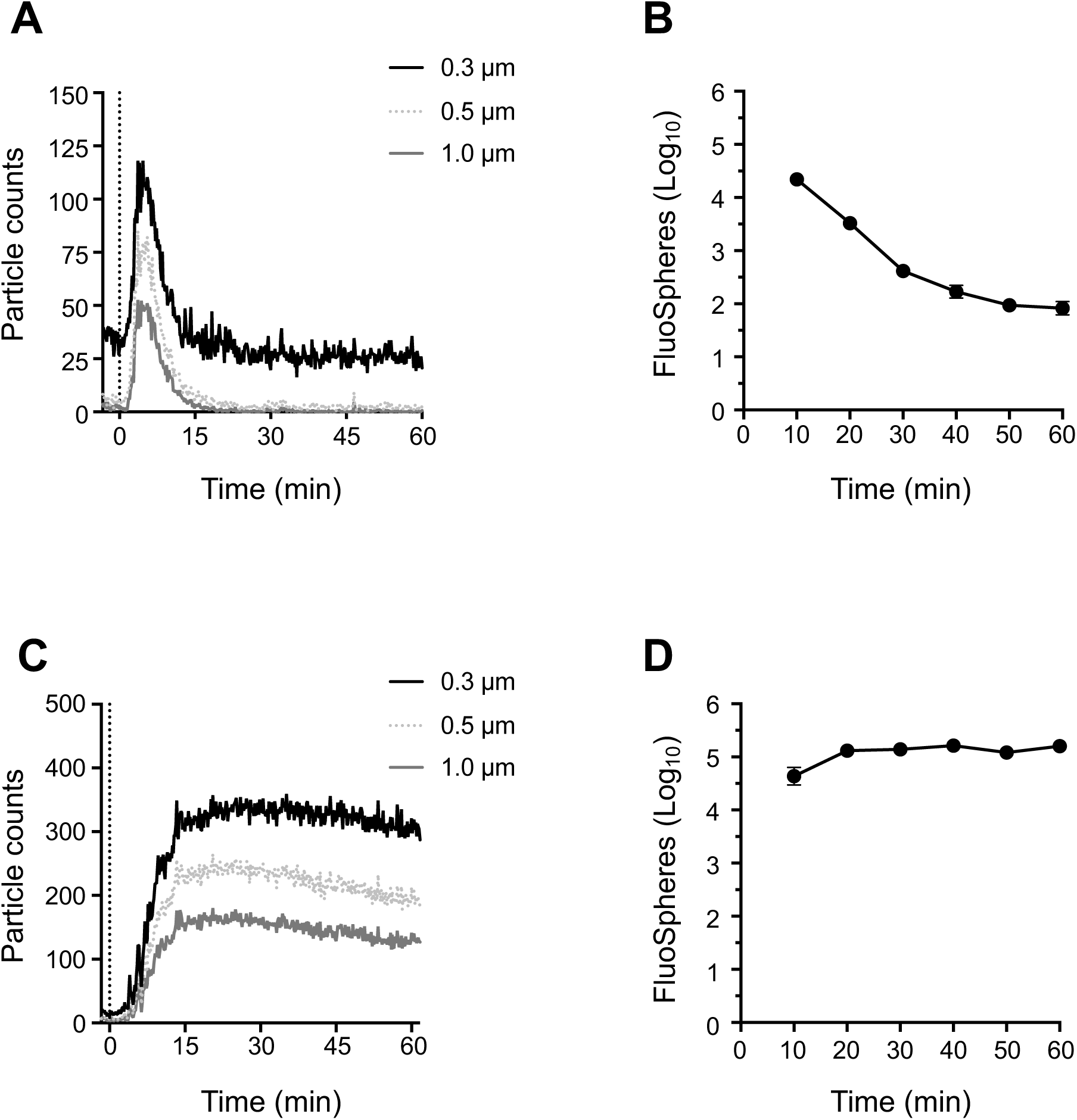
Aerosol release inside a BSL-3 laboratory. **A-D**, FluoSpheres (1 µm, 1×10^9^) were aerosolized inside a BSL-3 suite in the presence (**A-B**) or absence of forced ventilation (**C-D**). A Lighthouse particle counter was used to record airborne particles in the room. Coriolis air sampling was done with a HEPA-modified unit using the cumulative sampling method. **A** and **C**, Particle counts in the suite following aerosol dispersal. **B** and **D**, Detection of airborne FluoSpheres on the Coriolis determined by flow cytometry. Data from 3 experimental repeats shown. Error bars depict standard error of the mean. Dashed line indicates time of aerosol dispersal.

## DISCUSSION

The first line in protective measures against microbiological airborne threats is the ability to detect the pathogen in air, as it enables the mounting of adequate countermeasures in the next step, such as treatment, containment or disinfection, which greatly limits human exposure, prevent illness and save lives. This requires microbiological air-sampling tools that can be used in conjunction with analysis methods to rapidly detect microbes in air. In this regard, portable, tactical collectors are particularly useful in infection control as they can be widely distributed throughout society for air surveillance and research purposes. The Coriolis^®^ µ is a commercially-available, fieldable solution for air sampling that uses cyclonic technology to collect airborne particles directly into 15 mL buffer. Our study presents operative and technical modifications to the Coriolis to circumvent caveats during sampling and to improve its deployment. Building on the increasing number of applications for this collector, we reveal its suitability in collecting aerosols generated through simulated spills or experimental aerosolization in a BSL-3 laboratory.

The capacity of an air sampler to pull and precipitate airborne particles onto its collector piece is a bottleneck in the sampling process. In addition, since the amount of a target pathogen in air is expected to be low, successful detection may require both continuous monitoring and high-volume air sampling. Knowing an air sampler’s particle collection efficiency (Ladhani et al., 2017; Kim et al., 2018), may help identify the sampling conditions needed for successful collection, but useful, *bona fide* metrics for this measurement are difficult to obtain. Important previous work has generated measurements of relative sampling efficiency for several air samplers by benchmarking collection against the SKC BioSampler, showing that the Coriolis performed equally well against several aerosol-test agents including *Bg* and fluorescent microspheres (Dybwad et al., 2014). In our investigation of Coriolis performance, we evaluated the rate at which airborne microparticles were removed from the air during active sampling and report an effective clearance rate of approximately 35 L_air_/min. This is the rate at which the Coriolis clears 1 μm particles from air and not a direct measure of particle collection on the device. Still, this value could be used to estimate operation time in a given volume.

Coriolis collection greatly suffered from the harsh nature of cyclonic sampling as we observed sample loss and re-aerosolization from the collector cone. Both could contribute to misleading analysis results. Re-aerosolization was responsible at least in part for disseminating collected particles to various other parts of the sampler. This could inadvertently expose the user to pathogens during microbiological air sampling. Introduction of a cumulative sampling protocol and a HEPA-filter adaptation to the device improved these shortcomings. We have recently used the HEPA-modified Coriolis with the cumulative sampling protocol in our aerosol chamber to investigate the performance of a portable electrostatic air sampler for tuberculosis (Rufino de Sousa et al., 2020). A similar HEPA-modified Coriolis has also been used in a clinical, experimental setting to quantify *M. tuberculosis* from human bioaerosols (Patterson et al., 2017). In the current study we provide a thorough presentation of these technical and operational improvements to the Coriolis and supply details for assembly of the HEPA filter so that others can benefit from this adaptation. The HEPA-modified Coriolis operates otherwise like the standard, commercially-available unit. We observed a small increase in Coriolis particle clearance rate upon mounting the HEPA filter. This is probably due to the HEPA filter trapping airborne particles that would otherwise be subject to continuous re-circulation through the device during operation.

Despite many safety precautions and protective measures, handling live pathogens in the laboratory is a standing risk for occupational exposure, even to the most experienced staff. We thought it interesting to employ the Coriolis with improvements in the assessment of simulated incidents in the laboratory coupled to microbiological exposure. Here, collectors such as the Coriolis may bring important insight on exposure that may impact on future biosafety regulations and recommendations. In this context, following a substantial microbiological spill, it is generally recommended by biosafety delegates that personnel should vacate the room for 20-30 minutes due to the risk of exposure to aerosols (WHO, 2004). Cleaning and decontamination procedures are consequently delayed although robust experimental support for this risk assessment is missing. Using the Coriolis and particle counters, we show that a simulated spill with a large, concentrated volume of microspheres does not generate a significant number of aerosol particles in the environment. Because few airborne particles were generated in the spill, it might not have been a preferred simulation to study Coriolis performance. Nonetheless, it gave insight into an important and common biosafety issue related to microbiological exposure in the laboratory work environment. Our data thus suggests that infection control measures can be applied immediately after a large microbiological spill since the risk of aerosol dissemination and exposure to the user in this condition is negligible. It is unclear to what degree dust particulates from a surface could be re-aerosolized during a spill to potentiate airborne particles and microbial exposure. *B. anthracis* spores have been reported to be re-aerosolized from contaminated office surfaces under conditions of low personnel activity (Weis et al., 2002). A review of historical data on tuberculosis transmission highlights the risk of dust-borne *M. tuberculosis* in spreading the infection (Martinez et al., 2019). A premise for our recommendation of immediate decontamination is therefore that it be performed on laboratory surfaces that are otherwise kept clean and accumulation of dust minimized.

In the unique setting of our BSL-3 infrastructure with forced ventilation, we also used the Coriolis to investigate detection of aerosols carrying microspheres aerosolized on a BLAM, a Collison-type nebulizer. Collison nebulizers are readily used to experimentally infect laboratory animals through the aerosol route (May, 1973; Roy and Pitt, 2012). This experiment thus simulates a potential incident in an animal BSL-3 or aerobiology laboratory with ensuing infectious bioaerosol dispersal. Even though forced ventilation returned particle counts in the room to background levels within 15 min after the simulated incident, microspheres could be detected with the aid of the Coriolis up to 1 hr after aerosol release. In the absence of forced ventilation, particle numbers remained high and elevated for the entire duration of the experiment. These experiments reveal the importance of ventilation in limiting transmission of infectious bioaerosols. They also indicate that the risk of exposure remains for at least 1 hr after a *bona fine* aerosolization, even in the presence of forced ventilation. Thus, our simulations show that an accident with an aerosol generator introduces a much higher risk for occupational exposure compared to a large (0.5 L) microbiological spill.

## CONCLUSION

The field of aerobiology has been hampered by the lack of tactical (fieldable) units for microbiological air-sampling. Units such as the Coriolis are helping to fill this gap by providing a useful tool for the study and quantification of infectious bioaerosols. Our simple operative and technical modifications to the Coriolis should add to its biosafe deployment and promote continued investigation on human transmission and exposure to airborne pathogens.

## Acknowledgments

We thank Roland Möllby (Karolinska Institutet, Sweden) and Wayne Bryden (Zeteo Tech, USA) for suggestions and critical reading of this manuscript. We would also like to thank Sören Hartmann and Per-Erik Björk (Karolinska Institutet) and Erik Ekstedt (Akademiska Hus AB, Stockholm, Sweden) for technical assistance. Flow cytometry was performed at the Biomedicum Flow Cytometry Core facility (BFC), Department of Microbiology, Tumor and Cell Biology, Biomedicum, Karolinska Institutet.

---

**Supplementary Figure 1.**
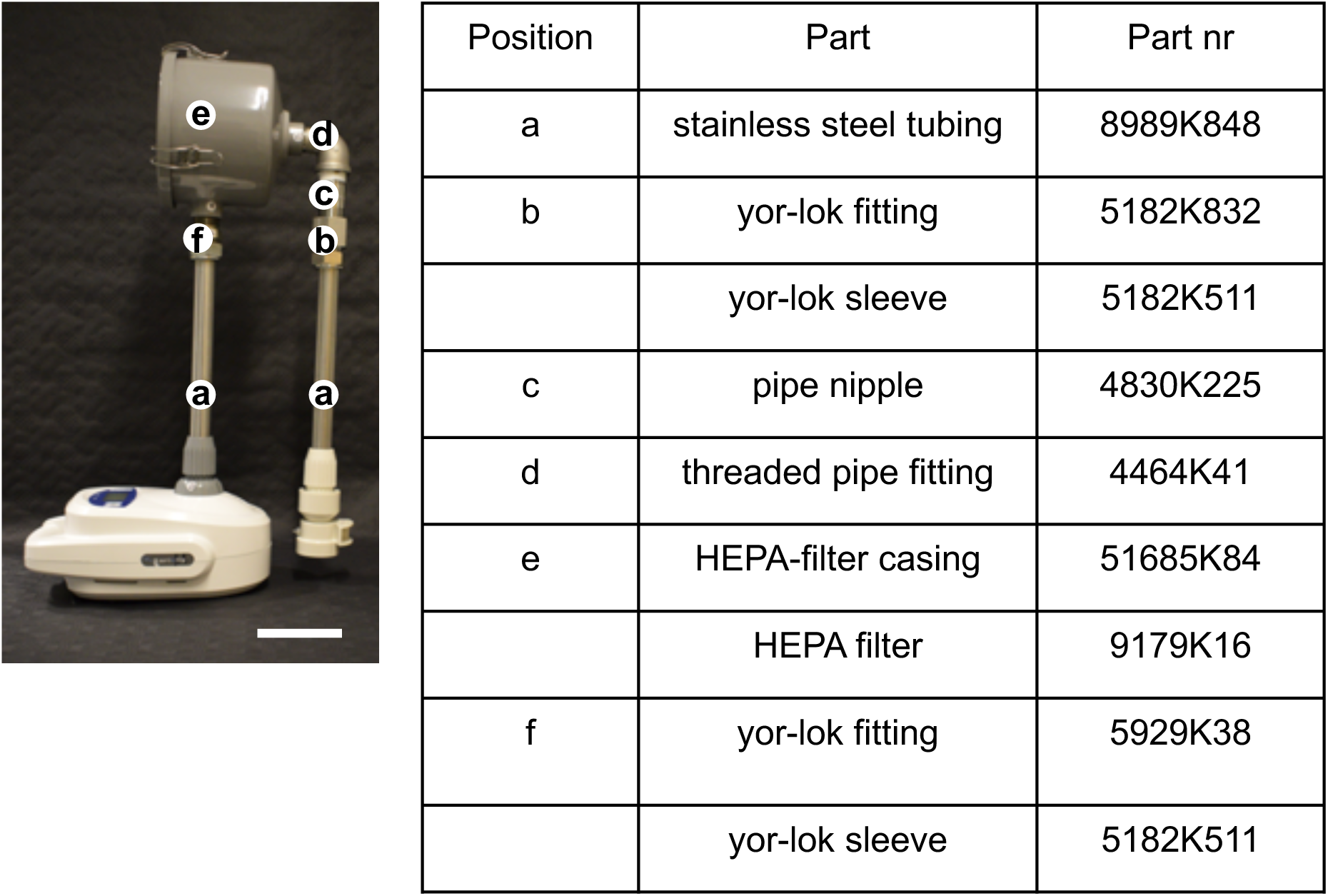
Parts description for Coriolis HEPA adaptation. Photograph and parts description including catalog number (all MacMaster-Carr, USA) for assembly of HEPA adaptation, amounting in all to about 745 USD (2020). Scale bar depicting 10 cm.

